# Detection of a novel enterotropic *Mycoplasma gallisepticum*-like in European Starling (*Sturnus vulgaris*) around poultry farms in France

**DOI:** 10.1101/2021.06.18.448922

**Authors:** Chloé Le Gall-Ladevèze, Laurent-Xavier Nouvel, Marie Souvestre, Guillaume Croville, Marie-Claude Hygonenq, Jean-Luc Guérin, Guillaume Le Loc’h

## Abstract

Recent outbreaks of highly pathogenic avian influenza in southwest France have raised questions regarding the role of commensal wild birds in the introduction and dissemination of pathogens between poultry farms. To assess possible infectious contacts at the wild-domestic bird interface, the presence of *Mycoplasma gallisepticum* (MG) was studied in the two sympatric compartments in southwest France. Among various peridomestic wild birds (n=385), standard PCR primers targeting the 16S rRNA of MG showed a high apparent prevalence (up to 45%) in cloacal swabs of European starlings (*Sturnus vulgaris*, n=108), while the MG-specific *mgc2* gene was not detected. No tracheal swab of these birds tested positive, and no clinical sign was observed in positive birds, suggesting commensalism in the digestive tract of starlings. A mycoplasma strain was then isolated from a starling swab and its whole genome was sequenced using both Illumina and Nanopore technologies. Phylogenetic analysis showed that it was closely related to MG and *M. tullyi*, although it was a distinct species.

A pair of specific PCR primers targeting the *mgc2*-like gene of this MG-like strain was designed and used to screen again the same avian populations and a wintering urban population of starlings (n=50). Previous PCR results obtained in starlings were confirmed to be mostly due to this strain (20/22 positive pools). In contrast, the strain was not detected in fresh faeces of urban starlings. Furthermore, it was detected in one tracheal pool of cattle egrets and one cloacal pool of white wagtails, suggesting infectious transmissions between synanthropic birds with similar feeding behaviour. As the new starling mycoplasma was not detected in free-range ducks (n=80) in close contact with positive starlings, nor in backyard (n=320) and free-range commercial (n=720) chickens of the area, it might not infect poultry. However, it could be involved in mycoplasma gene transfer in such multi-species contexts.

## 1. INTRODUCTION

In a context of increasing global concerns about animal welfare, free-range livestock farming is on the rise in industrialised countries (Sørensen, Edwards, Noordhuizen, & Gunnarsson, 2006; Thornton, 2010), including in the poultry production sector (Bessei, 2018). Due to the nature of free-range systems, an increase in free-range farming mechanically leads to an increase in the farmland devoted to production. This leads in turn to larger areas where farm animals and wildlife may come into contact. As livestock spend more time outdoors, free-range farming also increases the probability of contacts between livestock and wildlife. Depending on environmental conditions and the species present on both sides of this interface, micro-organisms can be shared via direct and indirect contacts. This enriched microbiome is composed predominantly of commensal strains that could be considered to be protective flora, but some of these shared opportunistic infectious agents can occasionally be or become pathogenic (Bass, Stentiford, Wang, Koskella, & Tyler, 2019; Vayssier-Taussat et al., 2014).

Among the infectious agents that can be shared between species, mycoplasma show a wide variety of strains, the majority of which are commensal and non-pathogenic (Citti & Blanchard, 2013). However, some can cause pathogenicity in their hosts. These include *Mycoplasma gallisepticum* (MG), which is the most frequent mycoplasma pathogen of poultry and causes chronic respiratory syndromes with considerable economic and sanitary impacts (Ferguson-Noel, Armour, Noormohammadi, El-Gazzar, & Bradbury, 2020; Levisohn & Kleven, 2000). Because domestic Galliformes are its main natural reservoir, the detection of MG in wild birds living in contact with poultry farms is likely an indicator of a spillover from livestock to wildlife. In eastern USA, such a spillover was suspected to have occurred in House Finch (*Haemorhous mexicanus*), and the species was found to be a secondary maintenance host (Fischer, Stallknecht, Luttrell, Dhondt, & Converse, 1997; Luttrell et al., 2001). The spread in commensal wild birds generated fears of possible spillbacks to commercial poultry as the transmission of the bacteria via direct contacts and materials was established (Stallknecht, Luttrell, Fischer, & Kleven, 1998; Luttrell et al., 2001; Dhondt, Dhondt, & Ley, 2007), and because an atypical strain isolated in commercial turkeys was genetically related to house finch strains (Ferguson, Hermes, Leiting, & Kleven, 2003). Natural infections of wild birds with MG are reported worldwide, and have been described in various taxa (Dhondt, DeCoste, Ley, & Hochachka, 2014; Ferguson-Noel et al., 2020; Ley, Hawley, Geary, & Dhondt, 2016; Michiels et al., 2016; Sawicka, Durkalec, Tomczyk, & Kursa, 2020; Sumithra et al., 2013). This wide potential of distribution associated with various transmission modes and consistent environmental survival suggest that the possibility of spillovers and spillbacks of MG between domestic and wild birds is significant.

MG in field samples is conventionally identified using bacterial cultures with immunological or PCR methods, or directly by PCR detection on the original sample. As mycoplasma cultivation is often long and fastidious, PCR methods generally are preferred when rapid results are needed, such as in routine clinical diagnosis or large epidemiological studies. The usual MG DNA sequence targets are 16S rRNA (Lauerman, 1998), defined as a reference target by OIE (OIE, 2019), and multi-locus sequence typing (MLST) sequences such as the *mgc2* gene (García, Ikuta, Levisohn, & Kleven, 2005). The latter was defined as a more specific target, in contrast with PCR primers for the highly conserved 16S rRNA sequence that might amplify other MG-like species hosted by birds such as *Mycoplasma imitans* (García et al., 2005), *Mycoplasma tullyi* (Yavari et al., 2017), and other unidentified species detected in seabirds (Ramírez, 2019).

To assess possible infectious contacts at the wild-domestic bird interface, the presence of MG was studied in the two sympatric compartments in southwest France. The region is of particular interest for studying the wild-domestic bird interface as it is the second largest poultry production area in France, with various species (Galliformes and Anseriformes) and a high density of quality labels and foie gras production units that require poultry to be kept outdoors for several weeks. These conditions offer frequent potential occasions for contacts between wild and domestic compartments, which has led, for example, to investigations into the role of wild birds in the diffusion of Highly Pathogenic Avian Influenza Viruses (HPAIV) during the last epizootics of 2015-2016, 2016-2017 and 2020-2021 (Anses, 2016; Guinat et al., 2017; Van De Wiele et al., 2017; Plateforme ESA, 2021). As MG may rely on the same direct and indirect transmission routes as influenza viruses, with longer expected excretion times in its host, it could be used as a transmission marker between wild and domestic birds in this region. This study aims to investigate this hypothesis and characterize the epidemiology and phylogeny of detected MG strains among several species of commensal wild birds of poultry farms, as well as among free-range commercial and backyard poultry flocks in the same area.

## 2. MATERIALS AND METHODS

### 2.1. Sampling birds at the wild-domestic interface

#### 2.1.1. Sampling of wild birds

Samples were taken in two separate campaigns. The first took place between December 2016 and February 2017 in the framework of highly pathogenic avian influenza surveillance in wild birds under the authority of the prefecture and government wildlife services. Captures were performed by lethal shooting of healthy birds living in close proximity to poultry farms in the departments of Gers and Hautes-Pyrénées in southwestern France. Sampled birds consisted of 89 European starlings (*Sturnus vulgaris*) and 29 cattle egrets (*Bubulcus ibis*) (Table 1). Both oropharyngeal and cloacal swabs were taken from each individual. All samples were stored in 300μL sterile 1% PBS on ice before being frozen at −80°C in the laboratory.

**Table 1.**
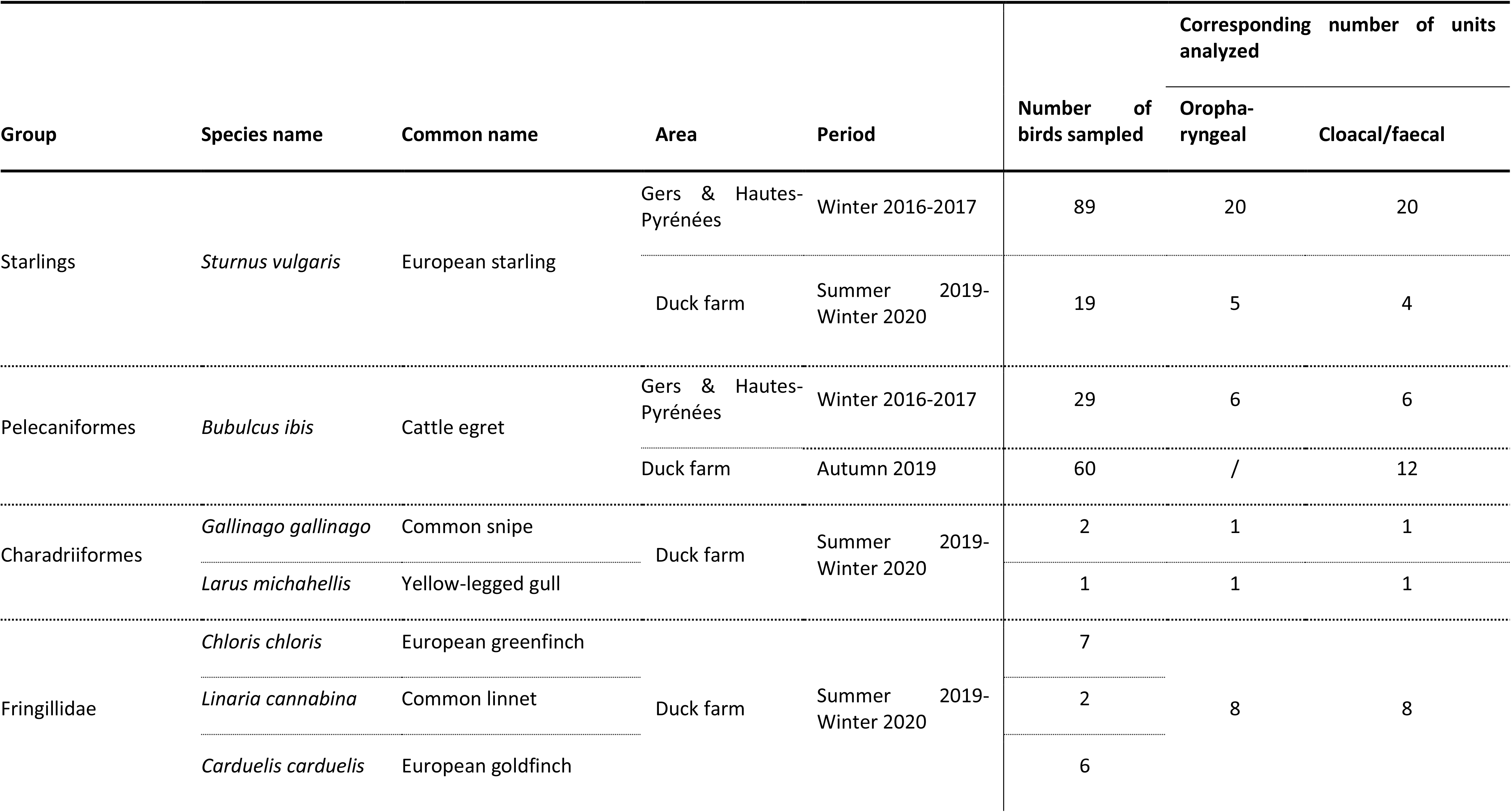
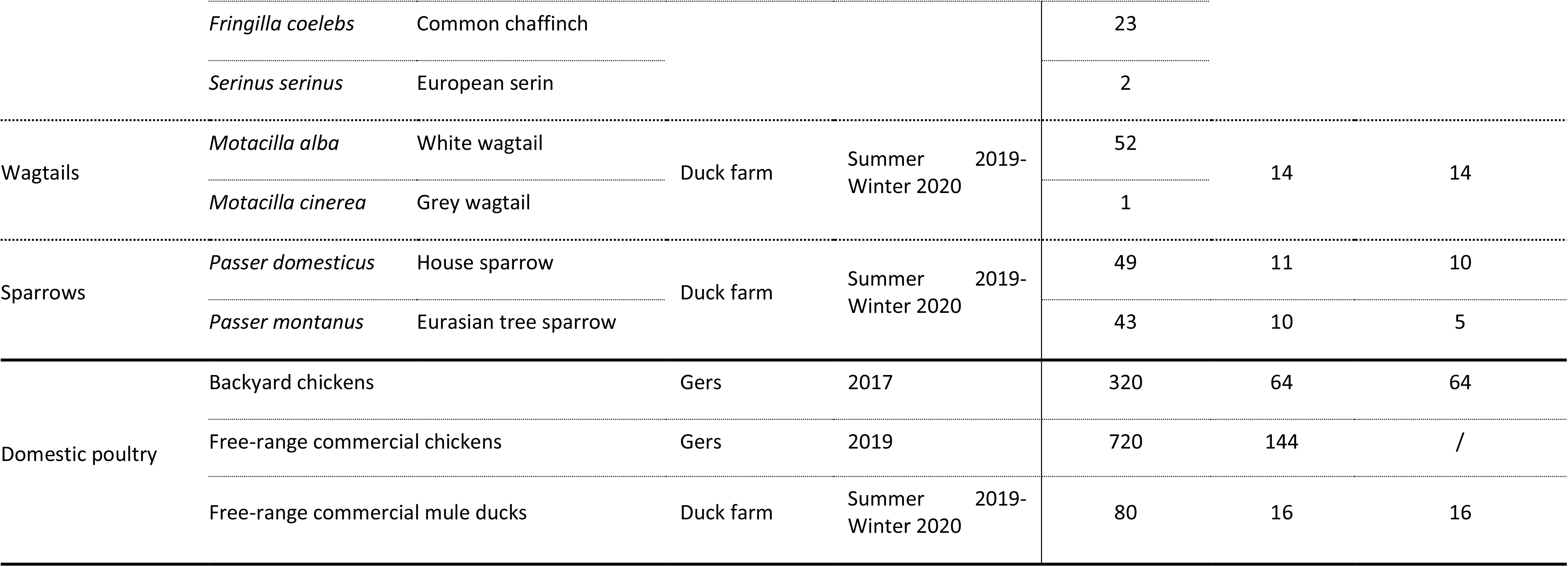
Description of wild and domestic bird groups sampled, and the corresponding number of sample pools analyzed for each group.

The second sampling campaign took place from July 2019 to February 2020. It was based on mist-net captures targeting healthy small to medium-sized wild species on a mule duck farm in the department of Gers. Sampled birds consisted of 207 individuals representing 12 species, including 40 birds of five species in the *Fringillidae* family, 92 sparrows (*Passer domesticus* and *Passer montanus*), 53 wagtails (*Motacilla alba* and *Motacilla cinerea*), 19 European starlings, and three Charadriiformes (Table 1).

Again, both oropharyngeal and cloacal swabs were taken from each individual. All wild bird captures and manipulations were performed with the help of a professional bird-bander, and the protocol was subjected to a mandatory authorization by the Muséum National d’Histoire Naturelle, Paris, France. Samples were stored as previously described.

In parallel with the captures, in October and November 2019 a non-invasive method was used to sample a group of cattle egrets seen foraging during the autumn and winter months in close contact with farmed ducks and other wild species. A previously disinfected plastic surface was placed under their night roosts, which were located 2 km away from the farm, and 30 fresh faeces were collected using sterile swabs on one morning of each month. Moreover, in December 2019, a population of starlings spending winter nights in the centre of Toulouse city (around 100 km from the duck farm in Gers) was sampled in the same non-invasive way. Fifty fresh faeces were collected using sterile swabs on one morning under roosting trees used by starlings only. Samples were stored as previously described.

#### 2.1.2. Sampling of domestic birds

In 2017, in the same context of influenza investigations, 320 apparently healthy chickens were sampled from 64 backyard flocks close to farm outbreaks in the same department of Gers (Table 1). Tracheal and cloacal swabs were collected on five birds in each backyard flock and stored in 300μL sterile 1% PBS on ice before being frozen at −80°C.

In 2019, 720 commercial free-range chickens aged 84 days and originating from 36 farms were sampled post-mortem in slaughterhouses in Gers (Table 1). Tracheal swabs were collected following the standard regulatory sampling for MG diagnosis. Samples were stored as previously described.

In parallel with the wild bird captures in 2019-2020, mule ducks from the farm in Gers were sampled: tracheal and cloacal swabs from 20 ducks of the same flock, around 10 weeks of age (i.e., after 7 to 9 weeks of free-ranging life) were collected in April, November, December 2019 and February 2020 (Table 1). Samples were stored as previously described.

### 2.2. Detection and identification of mycoplasma strains

#### 2.2.1. Nucleic acid extractions

Because the swabs were subjected to screening for several viral and bacterial agents, RNA and DNA extraction were performed using QIAamp Viral RNA Mini Kit (QIAgen®) with spin-columns, following the manufacturer’s instructions. For extraction and subsequent analyses, all of the samples were pooled (five samples in each pool). Negative extraction controls were processed alongside each series of extractions. All extracted nucleic acids were then stored frozen until further analysis.

#### 2.2.2. Molecular screening

PCR screenings targeting 16S rRNA, *mgc2* adhesin and *mgc2*-like adhesin (list of primers used in Table 2) were made by real-time PCR (rtPCR) on LightCycler 96 thermocycler (Roche®). The LightCycler SYBR Green I Master reaction mix (Roche®) was used with the following volumes for each sample analyzed: total volume of 20μL, with 2μL of DNA sample, 0.4μL of each 10μM primer (final concentration of 0.2μM), 10μL of SYBR Green I Master, 7.2μL of PCR-grade water. Negative and positive PCR controls consisted respectively of 2μL of water and 2μL of standard dilutions of plasmid construct including the specific PCR target sequence. All thermocycling programs were then set on the same following conditions: pre-incubation step of 5min at 95°C, 45 amplification cycles of 10s at 95°C, 15s at 55°C and 15s at 72°C with single acquisition, and a melting analysis phase of 10s at 95°C, 1min at 65°C and 1s at 97°C with continuous acquisition. Positive results were considered as showing a cycle threshold (Ct) under 44 and a melting temperature peak in a 2°C range around the one of positive controls.

**Table 2.**
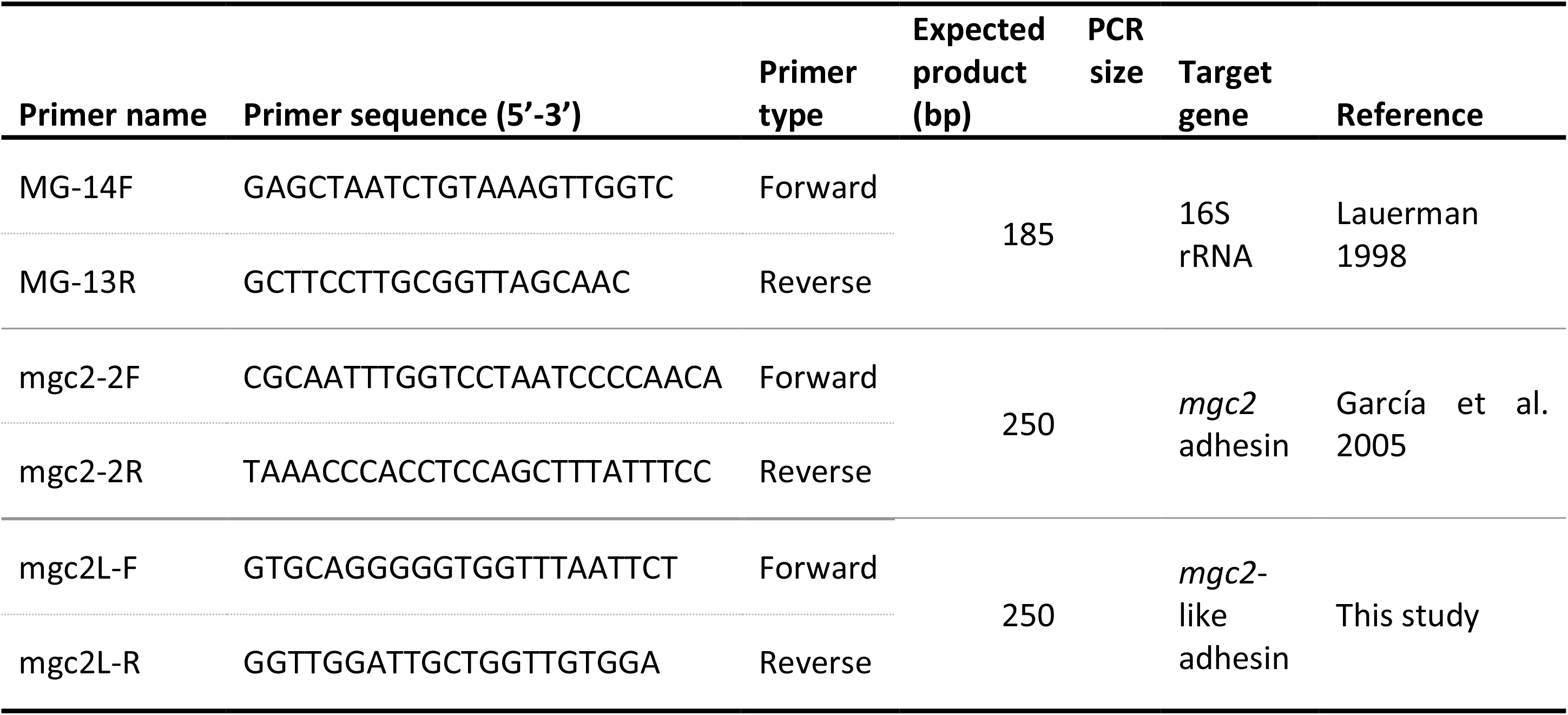
Description of PCR primers used in this study

#### 2.2.3. Cultivation and isolation of mycoplasma strains

Mycoplasma were cultured from the most loaded starling cloacal swabs in sequential incubations and purifications following standard protocols (Tully, 1995) adapted for contaminated samples. Briefly, swabs were initially enriched by incubation in 1.8mL SP4 liquid growth medium supplemented with Thallium acetate (0.1% w/v) for 4h at 37°C. First 0.1 dilutions of these broths were plated on agar, but resulted in non-mycoplasmal contamination covering colonies with a typical fried-egg shape. To purify and select the most of the target mycoplasma from the first enrichment broth, two supplementary passages were realised, each one cultured at 37°C for 3 days until the colour of the medium changed (to bright yellow). The entire volume of the resulting enriched broth was then filtered at 0.45μm. As no growth resulted on agar plates from the filtrate, it was subjected to a supplementary passage and cultured in a liquid medium at 37°C for 3 days until the colour changed. Plating on agar of 10μL from this last filtered-enriched broth resulted in one single mycoplasma colony from one of the initial swabs. The identity of the isolated strain of Starling mycoplasma was confirmed by PCR, then it was enriched and kept frozen for further use.

### 2.3. Phylogenetic and epidemiological characterization of the novel Starling mycoplasma

#### 2.3.1. Whole genome sequencing

Genomic DNA was extracted and purified from a broth culture of Starling mycoplasma, using a standard phenol-chloroform method (Chen & Kuo, 1993). The DNA then was sequenced using two complementary sequencing chemistries: Oxford Nanopore Technologies (ONT) and Illumina Hi-Seq. The ONT library was prepared using the field sequencing kit (SQK-LRK001) and loaded on a flongle flowcell (FLO-FLG001), then the sequencing run was started on a MinION MK1B device for 10 hours. Nanopore raw reads were basecalled using Guppy (v2.2.3), and a first assembly was performed using Canu (v1.6) to assess the possibility of constructing a draft genome based on nanopore reads. The DNA also was sequenced by Illumina Hi-Seq (paired-end, 2×150 bp), performed at the GATC Biotech facility (Konstanz, Germany). A hybrid assembly combining ONT and Illumina reads was made using Unicycler version 0.3.4 (Wick, Judd, Gorrie, & Holt, 2017). The resulting whole genome scaffold was annotated with NCBI prokaryotic genome annotation pipeline (Tatusova et al., 2016). The nucleotide sequence is accessible on GenBank under accession number CP068418.

#### 2.3.2. Phylogenetic analysis

Genome alignments, synteny and identity compared with reference strains of mycoplasma were determined and visualised using the wgVISTA tool (Frazer, Pachter, Poliakov, Rubin, & Dubchak, 2004).

Phylogenetic analyses were conducted using MEGA version X (Kumar, Stecher, Li, Knyaz, & Tamura, 2018) on the longest available 16S rRNA sequences. Reference strain sequences were retrieved from GenBank, aligned together with those of the Starling mycoplasma using MUSCLE algorithm, then compared by a maximum likelihood tree.

A search for antimicrobial resistance genes was performed using the CARD Resistance Gene Identifier version 5.1.1 tool (Alcock et al., 2020).

#### 2.3.3. Design of specific qPCR primers and protocol

Specific rtPCR for detection of the Starling mycoplasma was designed to target the domain coding for its *mgc2*-like adhesin. Selection of priming sites was made *in silico* using the Primer BLAST tool from NCBI (Ye et al., 2012), selecting a PCR-product size range of 150-250 bp, and excluding matches with the whole mycoplasma reference database. Among propositions of primer pairs, selection was based on the least self-complementarity of both primers. Selected primers *mgc2*L-F/*mgc2*L-R (Table 2, Appendix S2) were then tested *in vitro* to assess the best PCR conditions regarding sensitivity and specificity of amplification with genomic DNA of the new Starling mycoplasma, compared with *Mycoplasma gallisepticum* PG31, *M. gallinaceum* and *M. pullorum* strains. A plasmid construct including the PCR target sequence of the new mycoplasma was then built and used as a positive PCR control.

#### 2.3.4. Epidemiological analysis

As screenings were mainly performed on pooled samples, the estimated pooled prevalence and 95% confidence interval were calculated online with EpiTools (Sergeant, 2018) using the “Pooled prevalence for fixed pool size and perfect tests” tool with the method for exact confidence limits assuming a binomial distribution (Method 3 from (Cowling, Gardner, & Johnson, 1999)).

## 3. RESULTS

### 3.1. Distribution of MG and close relatives at the wild-domestic bird interface

Detection of MG DNA in wild bird and domestic poultry compartments gave distinct results (Table 3). In the wild compartment sampled in the winter of 2016-2017, 19 out of 20 cloacal pools from starlings (estimated pooled prevalence 45% [95% CI: 24-74], mean Ct 26.4) and 1 out of 6 oropharyngeal pools from cattle egrets (4% [95% CI: 0-19], Ct 43.3) were positive for MG 16S rRNA; however, none of them were positive for the *mgc2* gene. Results from the second sampling campaign in 2019-2020 revealed 3 out of 4 cloacal pools from starlings positive for MG 16S rRNA (24% [95% CI: 4-64], mean Ct 27.5), all negative for the *mgc2* gene, and no detection of MG 16S rRNA in 12 faecal pools from cattle egrets foraging on the same farm area. In other groups of wild birds sampled on the duck farm in 2019-2020, no Charadriiformes, Fringilliae or sparrows showed positive PCR results for MG 16S rRNA, while 1 out of 14 cloacal pools from wagtails was positive for MG 16S rRNA (1.5% [95% CI: 0-8], Ct 41.1) and negative for the *mgc2* gene. On the sampling campaign in urban starlings in 2019, all faecal pools were negative for MG and close relatives (Table 3).

**Table 3.**
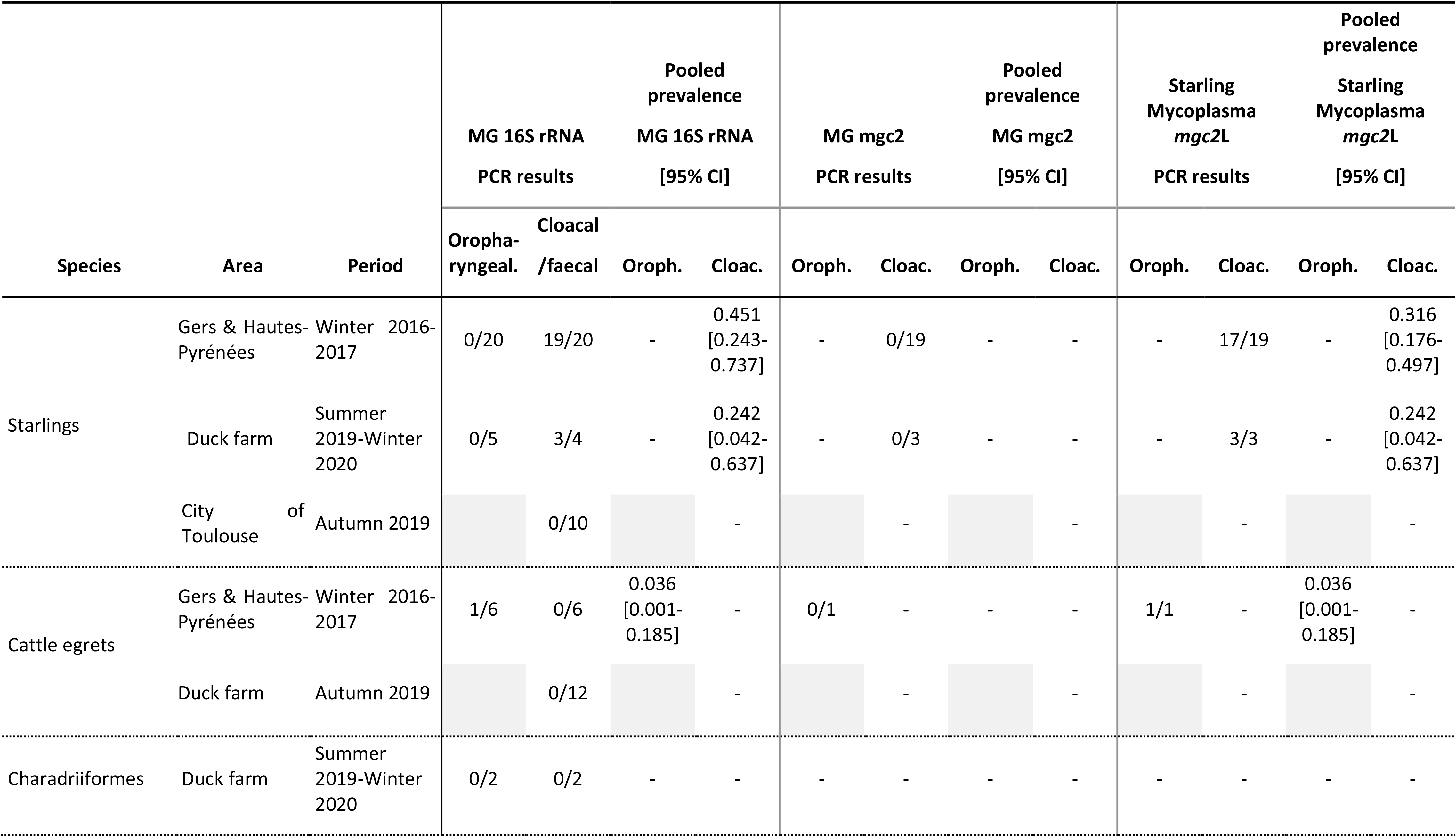
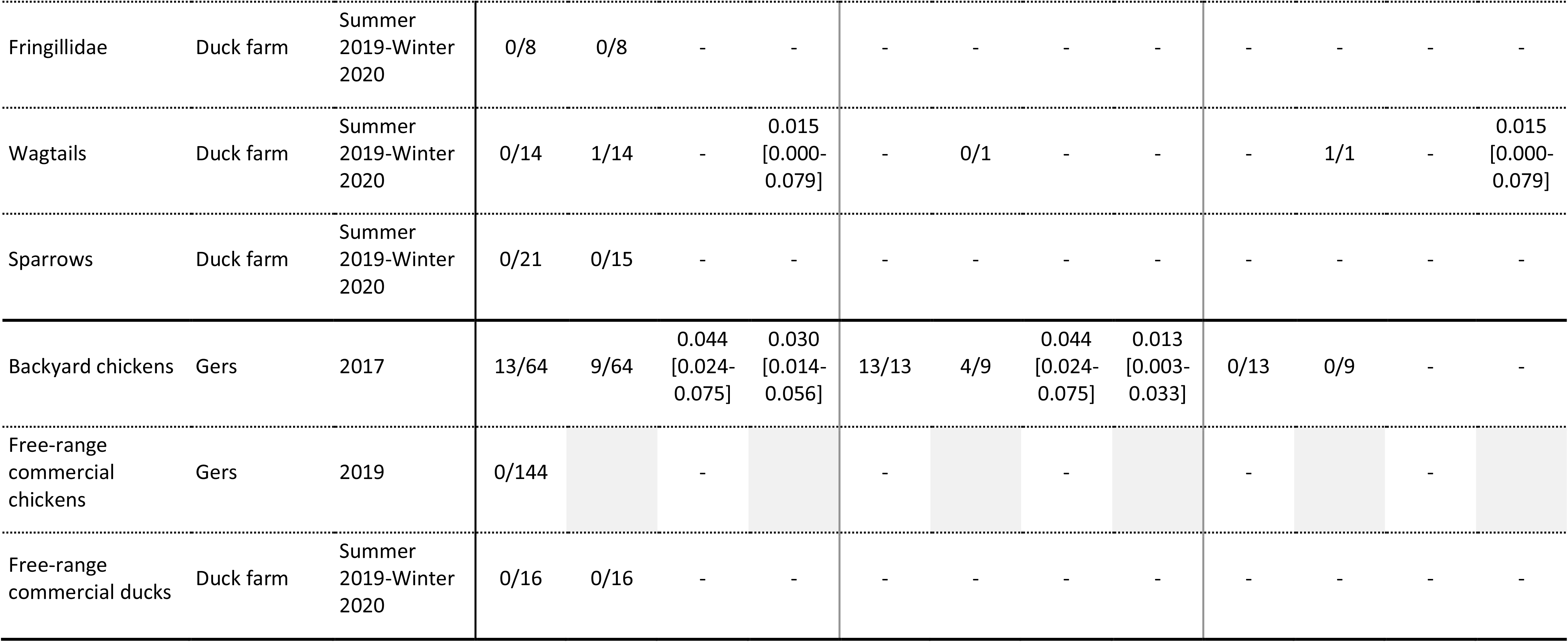
PCR results and corresponding pooled prevalence estimates for each group of wild and domestic birds sampled. PCR results are shown as number of positive pools or individuals over number tested. Oroph.: oropharyngeal samples. Cloac.: cloacal or faecal samples.

In domestic poultry, while none of the 144 commercial free-range chicken oropharyngeal pools were found positive for MG 16S rRNA, 13 out of 64 backyard chicken oropharyngeal pools tested positive for both MG 16S rRNA (mean Ct 40.3) and *mgc2* genes, corresponding to an estimated pooled prevalence of MG in backyard chicken oropharynx of 3% [95% CI: 1-6]. Regarding cloacal samples, MG 16S rRNA was detected in 9 out of 64 pools (mean Ct 39.4), followed by *mgc*2 confirmation in only 4 out of 9 pools, corresponding to an estimated pooled prevalence of MG in backyard chicken cloaca of 1% [95% CI: 0-3]. Ducks sampled on the farm in 2019-2020 all showed negative PCR results for MG 16S rRNA (Table 3).

### 3.2. Characterization and detection of a novel *Mycoplasma* species

As PCR results from wild birds showed a high prevalence for MG 16S rRNA in Eurasian starlings, cloacal swab supernatants from starlings showing the highest bacterial loads on PCR were cultured in mycoplasma medium. Colonies showed a typical fried-egg shape on agar plates, with very similar appearance to MG strains.

Whole genome sequencing from one of these colonies was carried out. Nanopore sequencing produced a total of 18,830 reads with a N50 value of 11.87 Kb. Canu analysis allowed the assembly of a draft genome in one contig of 938,090 bp. The Illumina sequencing produced 2×10,536,376 reads corresponding to an average of 3,400X for coverage depth. Hybrid assembly with Unicycler allowed one 925,667 bp circularized genome to be obtained.

Comparison of this genome to sequenced strains of MG, other close relatives found in domestic and wild birds and other mycoplasma species available in public databases revealed that this Starling mycoplasma isolated from cloaca belonged to a distinct new mycoplasma species. It was close to MG strain NCTC10115 (GenBank accession: LS991952, overall 91.1% identity, 16S rRNA 98.5% identity), *M. tullyi* isolated from a captive Humboldt penguin in UK (GenBank accession: CP059674, overall 91.1% identity, 16S rRNA 98.6% identity), and another unidentified mycoplasma detected in a wild Kelp Gull in South Africa (GenBank accession: FM878638, 16S rRNA 98.99% identity). However, this new species of Starling mycoplasma was very distinct from *M. sturni*, commonly isolated from the same host, with only 78.4% identity on its 16S rRNA sequence. The phylogenetic tree based on the whole 16S rRNA sequence confirmed these relations (Figure 1). The Starling mycoplasma genome showed highly similar regions with MG strain NCTC10115, such as its 16S rRNA sequence 98.5% identical (Appendix S1), or its 23S rRNA showing 97.5% identity. However, many regions were strongly different from MG homologous ones, in particular genes used for MLST of MG strains, such as *mgc2* sequence showing only 54.7% identity to its homologous one in the Starling mycoplasma (Appendix S2), or the 16S-23S rRNA intergenic region with 88.8% identity and the rpoB with 82% identity. Synteny visualisation of MG strain NCTC10115 and the Starling mycoplasma genomes shows conserved, repeated, or reversed homologous regions (Figure 2).

**Figure 1.**
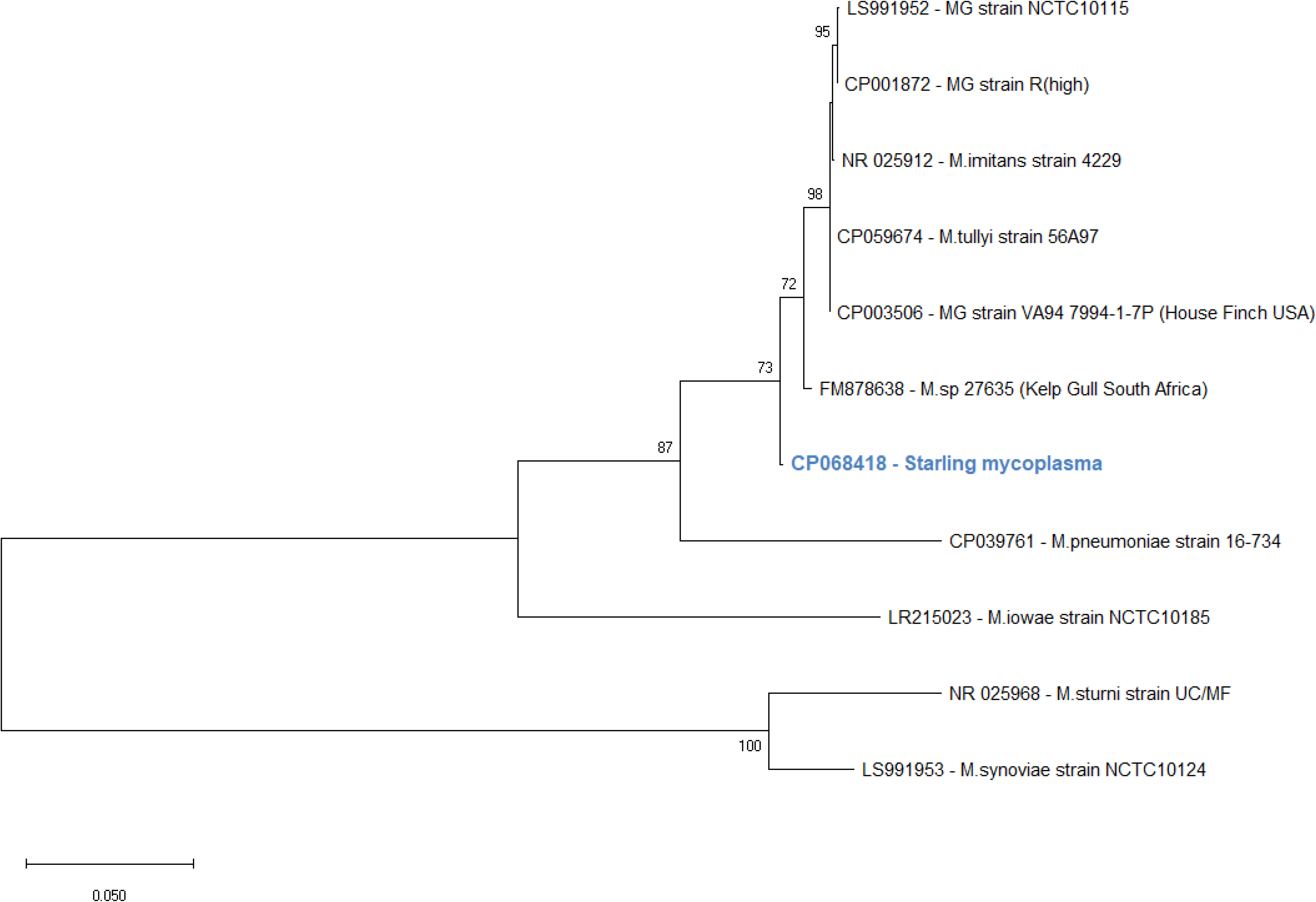
Phylogenetic tree based on 16S rRNA sequences of Starling Mycoplasma and various mycoplasma reference strains. The new Starling Mycoplasma is highlighted in blue. Minimum sequence alignment length is 1248nt. The maximum likelihood tree was built on a Tamura-Nei Gamma-distributed model, with 300 bootstrap replications. Bootstrap values lower than 70 are not shown

**Figure 2.**
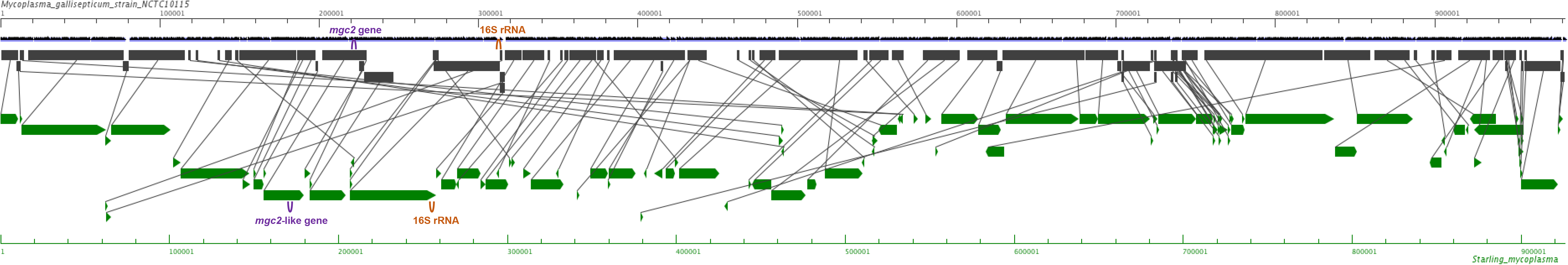
Whole genome synteny representation of Starling Mycoplasma and *Mycoplasma gallisepticum* strain NCTC10115 (LS991952.1). Loci for *mgc2* or “*mgc2*-like” adhesin (purple) and for 16S rRNA (orange) are indicated

No perfect hit was found in antimicrobial resistance genes of the Starling mycoplasma, with only 37 loose hits of 69% or less identity with partial Antibiotic Resistance Ontology reference genes.

### 3.3. Distribution of the novel *Mycoplasma sp*. at the wild-domestic bird interface

Using the specifically designed rtPCR for the *mgc2*-like gene, previous results from MG screenings were checked for presence of the novel Starling mycoplasma. In wild birds that were positive for MG 16S rRNA, the presence of the Starling mycoplasma DNA was confirmed in the oropharyngeal pool of 2016-2017 cattle egrets (3.6% [95% CI: 0.1-18.5], Ct 44.2), in 17 out of the 19 cloacal pools of 2016-2017 starlings (32% [95% CI: 18-50], mean Ct 34.1), in the three cloacal pools of 2019-2020 starlings (24% [95% CI: 4-64], mean Ct 39.9), and in the cloacal pool of wagtails (1.5% [95% CI: 0-7.9], Ct 39.1) (Table 3, Figure 3). Thus, only two of MG 16S rRNA positive pools of starlings were left unidentified.

**Figure 3.**
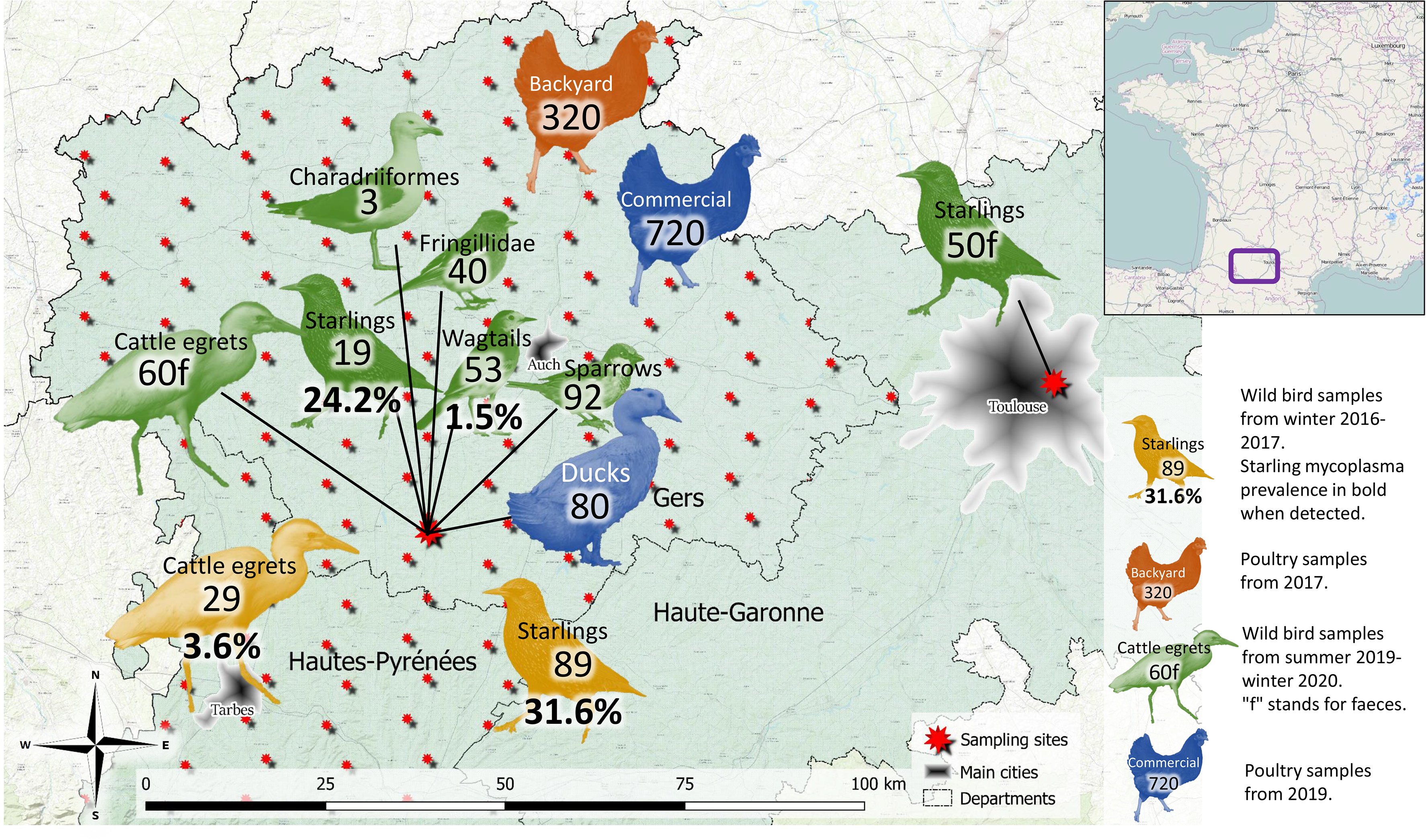
Map of Starling mycoplasma distribution in species, space and time

In the domestic compartment, the 13 oropharyngeal pools and 9 cloacal pools of backyard chickens that had been found positive for MG 16S rRNA showed no detection of the Starling mycoplasma (Table 3, Figure 3). As MG was previously detected in most of these pools, only 5 cloacal pools were left with no identification.

## 4. DISCUSSION

Looking for signs of infective contacts at the wild-domestic bird interface, a search for MG DNA was carried out in various commensal wild bird samples collected around poultry farms in the departments of Gers and Hautes-Pyrénées in southwest France. The widely used PCR test, based on standard MG14F/MG13R primers, was used as the first-line screening tool. As this PCR might be slightly unspecific (detection of MG and close relatives), a second PCR using *mgc2* primers was performed on positive samples in order to increase specificity for MG. Unexpected results were obtained when comparing both PCR. Indeed, while very few wild birds and backyard chickens carried 16S rRNA of MG, opposite results were obtained for European starlings around poultry farms. These starlings showed a particularly high apparent prevalence of MG 16S rRNA, but no detection of the *mgc2* gene, and a surprising strict cloacal tropism for bacteria usually considered as adapted to the respiratory tract. After isolation from one starling cloacal swab and whole genome sequencing of this atypical mycoplasma strain, comparative phylogeny with close species such as MG and *M. tullyi* revealed the identification of a new distinct species of mycoplasma. Screening using a specifically designed rtPCR for this novel mycoplasma confirmed its presence in the previous starling samples, but no DNA was detected in urban starlings of the same region. In the sympatric bird populations of positive starlings, no domestic poultry (chicken or ducks) were found to host the novel mycoplasma, and most of the other commensal wild birds also were found to be uninfected, except for one tracheal pool of cattle egrets and one cloacal pool of white wagtails.

Target species for captures and samplings were selected for their potential literature-based susceptibility to mycoplasmal infections (*Fringillidae*, Charadriiformes, starlings), or for their behaviour of high closeness with domestic birds (wagtails, sparrows, cattle egrets). In the USA, natural MG infections in house finches (*Haemorhous mexicanus*) have been detected for decades (Fischer et al., 1997; Luttrell et al., 2001), and it has been shown experimentally (Dhondt et al., 2007) and on field observations (Fischer et al., 1997; Ley et al., 2016) that other species of *Fringillidae* are also, but slightly less, susceptible. However, none of the 40 *Fringillidae* of our study were found to be infected by MG, even though some backyard poultry of the same region were found to be infected. Their role in the epidemiology of this bacteria in the studied area thus seems insignificant, contrary to what is observed in the USA. The other sampled wild species known to be susceptible to MG (Charadriiformes, starlings) did not seem to play a role in its regional epidemiology neither.

In contrast, the new mycoplasma was detected in a large proportion of starlings, and also in wagtails and cattle egrets. It is then interesting to note the presence of the new mycoplasma in species foraging in the same environment as infected starlings, which overall suggests infective contacts between these peridomestic species, and indicates the occurrence of occasional spillover events of this mycoplasma from its maintenance population (starlings) to other hosts. With only 3.6% estimated pooled prevalence in cattle egret respiratory tracts, and 1.5% in white wagtail digestive tracts, infection by the Starling mycoplasma seems to be more sporadic than endemic in these species. This idea is supported by the contrasting very high frequency of infection in sympatric starlings, which were excreting bacteria in their faeces and thus certainly infecting their shared foraging areas. It is also interesting to notice that this new mycoplasma can, at least sometimes, be found in the respiratory tract. In contrast, outdoor-ranging poultry did not show any contamination, although living in similar close conditions with infected starlings, supporting the idea that ducks and chickens might be less or not susceptible to the new mycoplasma. For the same reasons, Charadriiformes, *Fringillidae* and sparrows could be less susceptible to the Starling mycoplasma. In consequence, the bacteria may present a restricted natural host range in European starlings, with high prevalence levels in this population, up to 31.6%, and strict cloacal excretion, while very few spillover events occur to exposed sympatric avian species.

In order to compare the distribution of the Starling mycoplasma in different populations of its host, a wintering urban population of starlings was sampled in the same time range and approximately 100km away from the duck farm in Gers. None of the urban starling faeces collected were found to carry the mycoplasma, suggesting different epidemiological mechanisms. An explanation for the observed lack of detection could be the dilution of local possibly infected starlings into uninfected populations of starlings from eastern and northern Europe gathering during winter (Fliege, 1984; Feare & Douville de Franssu, 1992), a phenomenon that seems more important in urban areas (Peris, Motis, Martinez-Vilalta, & Ferrer, 1991; Robinson, Siriwardena, & Crick, 2005). A second and not exclusive hypothesis could be an absence of communication (i.e., different migration patterns) between rural and urban starling populations in the region related to different habitats and food availability (Feare & Douville de Franssu, 1992), combined with a restricted range of the starling mycoplasma to the rural starling population.

As the samplings targeted healthy individuals and no clinical signs were observed during manipulations or in the surroundings of capture fields, the novel Starling mycoplasma does not seem pathogenic to infected starlings, nor to spillover hosts like cattle egrets or white wagtails. Moreover, the cloacal tropism of the new mycoplasma seems atypical among other known related species such as MG in poultry, MG subspecies in house finches, *M. imitans* in various species, and *M. sp* 27635 in Kelp Gull (GenBank accession: FM878638 (Ramírez, 2019)), all isolated from respiratory tracts. However, mycoplasma are now known for their ability to horizontally transfer genes within and between species (Citti, Baranowski, Dordet-Frisoni, Faucher, & Nouvel, 2020; Citti, Dordet-Frisoni, Nouvel, Kuo, & Baranowski, 2018), giving the potential for a strain to jump in pathogenicity or infectivity to some host species on occasions of coinfection. The potential evolution of this new species in case of coinfection with closely related pathogenic strains then has to be considered, especially in this context of commensal avifauna in a region with a high density of free-ranging poultry and with both MG and Starling mycoplasma detected in coexisting populations.

Although a spillover of the Starling mycoplasma from wild birds to domestic poultry was not detected in the present study, and without regard to any genomic evolution of the mycoplasma, its circulation at the wildlife-livestock interface increases the chances of such transmission events to occur, with unknown clinical consequences in new hosts. Whether pathogenic or not, infection with the Starling mycoplasma could also be immunizing against close relatives such as MG. Further experimental infection studies are needed to assess the pathogenic and immunologic impacts of the infection by this species.

In addition to physiological effects, infection with the Starling mycoplasma in domestic birds could lead to diagnostic issues as highlighted by our results. In the poultry industry, diagnostic tests for MG are mostly made using PCR, the standard primer pair being MG14F/MG13R targeting the 16S rRNA gene (Jarquin, Schultz, Hanning, & Ricke, 2009; Lauerman, 1998; OIE, 2019). However, a lack of specificity has been described with these primers targeting a least variable region (Kempf, 1998), and alternative more MG-specific primers have been developed to tackle this issue (García et al., 2005; Raviv & Kleven, 2009). Among these new primers, the pair *mgc2*-2F/*mgc2*-2R targeting the adhesin *mgc2* gene seems to be the most reliable for rapid diagnosis thanks to high sensitivity and specificity for MG. While the needs for specificity are often low in indoor poultry production systems, and allow the use of 16S rRNA primers for MG detection, the complex multi-species context at the wild-domestic bird interface of our study pushes specificity needs to higher levels. Indeed, our results tackle a cross-testing issue when using 16S rRNA primers as a detection tool in contexts when atypical strains (pathogenic or non-pathogenic) or coinfections with MG can occur. In consequence, conventional detection with 16S rRNA PCR with no positive confirmation test in some research studies or pathology cases (e.g., Hartup & Kollias, 1999; Mikaelian, Ley, Claveau, Lemieux, & Bérubé, 2001) might incorrectly lead to a diagnosis of MG when some other closely related strain actually is the cause. Furthermore, occasional MG vaccination failures in domestic poultry could sometimes be due to infection with some atypical MG-like strain incorrectly diagnosed by the standard PCR test.

The present study describes the first complete genome of an MG-like species. This new data can help further phylogenetic studies in this group of poorly studied mycoplasma. In recent years, several other species of mycoplasma have been discovered in wild birds (Sumithra et al., 2013; Yavari et al., 2017; Oaks et al., 2004; Lierz, Hagen, Hernadez-Divers, & Hafez, 2008; Lecis et al., 2010) as molecular biology techniques have evolved to ease exploratory research. The strain described in this study is very close to *M. tullyi* isolated from a captive Humboldt penguin (Yavari et al., 2017), and to another unidentified mycoplasma *M. sp* 27635 detected in a Kelp gull from South Africa with only its 16S rRNA sequence published (GenBank accession: FM878638 (Ramírez, 2019)). Moreover, some cloacal samples of our study tested positive for MG 16S rRNA but negative with both *mgc2* or *mgc2*-like primers (2/19 starlings 2017, 5/9 backyards). This suggests the presence of other – maybe shared – unidentified strains or species of the “mycoplasmome” of bird cloaca that may cross-test with the standard 16S rRNA PCR, but also contribute in the gene pool transmitted horizontally between coinfecting mycoplasma strains.

## ACKNOWLEDGEMENTS

This study was performed in the framework of the “Chaire de Biosécurité Aviaire”, hosted by the National Veterinary College of Toulouse (ENVT) and funded by the Direction Générale de l’Alimentation, Ministère de l’Agriculture et de l’Alimentation, France. This project was also part of the AI-TRACK project, partly funded by the Occitanie Region and the French Comité Interprofessionnel des Palmipèdes à foie gras (CIFOG), France. ONT sequencing was performed in the framework of the FIELD project, in collaboration with the CIBU, Institut Pasteur, Paris, and funded by the France Futur Elevage Carnot Institute.

The authors wish to thank technical helpers without whom the study could not have been possible. In particular, Albert Phouratsamay (DVM) for sample collection on wild birds in the 2017 HPAI epizooty, Emma Hourantier (Master student) for PCR screening analyses on these wild bird samples, Adam Jbenyeni (DVM, PhD student) for helping in collecting poultry samples in Gers abattoirs, and Benjamin Vollot (licensed bird-bander for the Centre de Recherche sur la Biologie des Populations d’Oiseaux) for his efficiency in capturing and sampling wild birds on the duck farm. The authors also express their deepest gratitude to the entire Peres family, duck breeders in Gers, for their warm welcome at any time and for allowing us to capture birds around their farm.

## DATA AVAILABILITY STATEMENT

The data that support the findings of this study are available from the corresponding author upon reasonable request.

## ETHICAL APPROVAL

The authors confirm that the ethical policies of the journal, as noted on the journal’s author guidelines page, have been adhered to.

For sampling on live wild birds on the duck farm, CRBPO (Centre de Recherches sur la Biologie des Populations d’Oiseaux, Muséum National d’Histoire Naturelle of Paris) ethical review committee approval was received under the project authorization number 1035, and their guidelines for capturing and sampling wild birds were followed with the help of a professional bird-bander and following good veterinary practices. Live domestic poultry were handled and sampled under breeders’ authorization and following good veterinary practices.

## APPENDICES

## Appendix S1

## Appendix S2

