## Appendix S1 for "Detection of a novel enterotropic *Mycoplasma gallisepticum*-like in European Starling (*Sturnus vulgaris*) around poultry farms in France"

**Appendix S1** – 16S rRNA sequences alignment of Starling mycoplasma and MG strain R(high). Binding sites of conventional diagnostic primers for MG detection MG14F/MG13R (Lauerman, 1998) are framed in black.

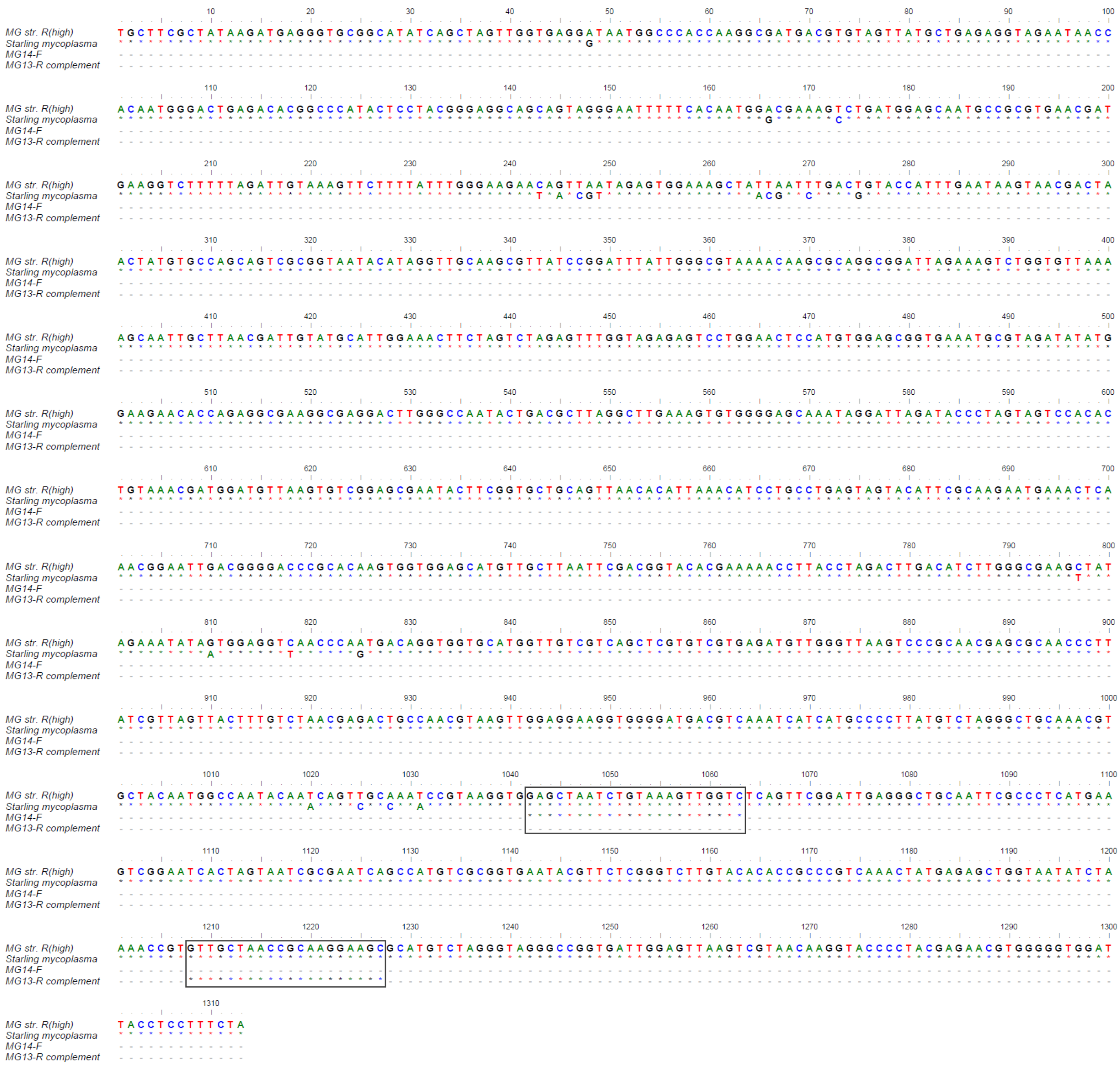
