## Appendix S2 for "Detection of a novel enterotropic *Mycoplasma gallisepticum*-like in European Starling (*Sturnus vulgaris*) around poultry farms in France"

**Appendix S2** – *mgc2* adhesin sequence of MG strain R(high) aligned with its homologous sequence "*mgc2*-like" on Starling mycoplasma genome. Binding sites of conventional diagnostic primers for MG specific confirmation mgc2-2-F/mgc2-2-R (García et al., 2005) are framed in black. Binding sites of specifically designed primers for Starling Mycoplasma detection are framed in bold purple.

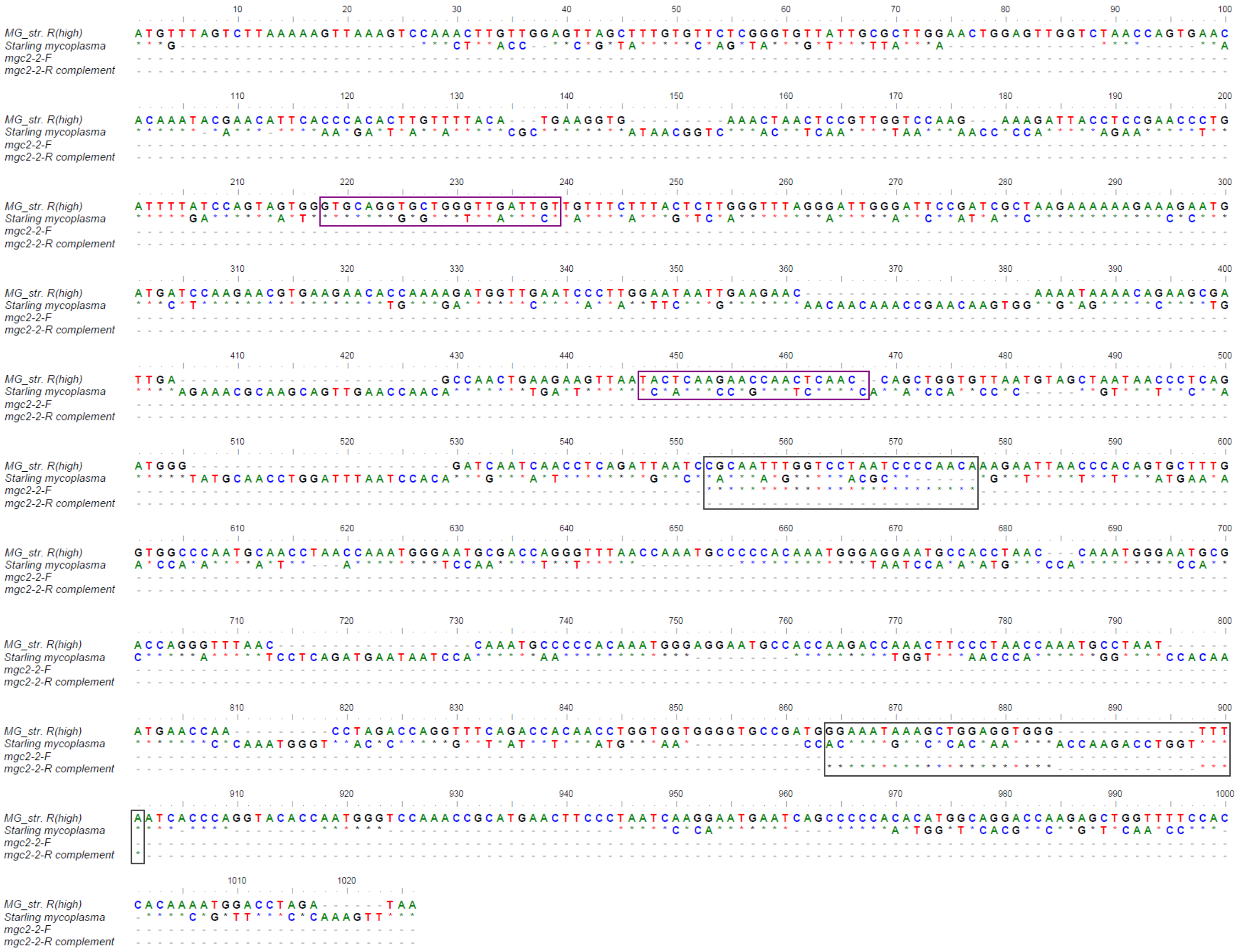
